# Endocannabinoid and neuroplasticity-related changes as susceptibility factors in a rat model of posttraumatic stress disorder

**DOI:** 10.1101/2024.03.06.583339

**Authors:** Laszlo Szente, Gyula Y Balla, Zoltan K Varga, Blanka Toth, Laszlo Biro, Zoltan Balogh, Matthew N Hill, Mate Toth, Eva Mikics, Mano Aliczki

## Abstract

Traumatic experiences result in the development of posttraumatic stress disorder (PTSD) in 10-25% of exposed individuals. While human clinical studies suggest that susceptibility is potentially linked to endocannabinoid (eCB) signaling, neurobiological PTSD susceptibility factors are poorly understood. Employing a rat model of PTSD, we characterized distinct resilient and susceptible subpopulations based on generalized fear, a core symptom of PTSD. In these groups, we assessed i.) eCB levels by mass spectrometry and expression variations of eCB system- and iii.) neuroplasticity-related genes by real-time quantitative PCR in the circuitry relevant in trauma-induced changes. Furthermore, employing supervised and semi-supervised machine learning based statistical analytical models, we assessed iv.) gene expression patterns with the most robust predictive power regarding PTSD susceptibility. According to our findings, in our model, generalized fear responses occurred with sufficient variability to characterize distinct resilient and susceptible subpopulations. Susceptible subjects showed lower prelimbic and higher ventral hippocampal levels of eCB 2-arachidonoyl-glycerol (2-AG) compared to resilient subjects. Ventral hippocampal 2-AG content positively correlated with the strength of fear generalization. Furthermore, susceptibility was associated with i.) neuronal prefrontal, hippocampal and amygdalar hypoactivity, ii.) marked decrease in the expression of genes of transcription factors modulating neuroplasticity and iii.) an altered expression pattern of eCB-related genes, including enzymes involved in eCB metabolism. Unsupervised and semi-supervised statistical approaches highlighted that hippocampal gene expression patterns possess strong predictive power regarding susceptibility. Taken together, the marked eCB and neuroplasticity changes in susceptible individuals associated with abnormal activity patterns in the fear circuitry possibly contribute to context coding deficits, resulting in generalized fear.

**Highlights:**

- We assessed endocannabinoid and neuroplasticity correlates of PTSD susceptibility
- Posttrauma generalized fear strength separates susceptible and resilient individuals
- Susceptible individuals show altered endocannabinoid levels in the fear circuitry
- Neuroplasticity and endocannabinoid gene expression patterns predict susceptibility

## 1. Introduction

Experiencing traumatic events can result in the development of posttraumatic stress disorder (PTSD), a severe psychiatric condition characterized by complex symptomatology, including re-experiencing the trauma, avoidance of trauma-related stimuli, emotional and arousal alterations, which symptoms last and fundamentally decrease life quality (American Psychiatric Association, 2022). In addition, trauma-associated learned fear often becomes generalized, i.e. fear responses are transferred to stimuli unrelated to the previous traumatic event, thus allowing fear responses to occur in safe situations as well (Lis et al., 2020; Sangha et al., 2020). As the lifetime prevalence of the disorder is relatively high (Kessler et al., 2005) and its therapy is currently insufficiently resolved (Davis et al., 2006), its pathophysiology is intensively studied with the ultimate goal of developing effective pharmacotherapeutic interventions. A crucial question in PTSD research is why only 10-25% of trauma-exposed individuals develop PTSD following a traumatic experience, while the majority of the population recover without long-term consequences (Breslau et al., 1998, 1991). Identification of the neurobiological basis of risk factors that increase susceptibility to developing PTSD following a traumatic event represents a major clinical challenge with high therapeutic potential.

The endocannabinoid (eCB) signaling system – consisting of cannabinoid receptors. eCBs 2-arachidonoylglycerol (2-AG) and N-arachidonoylethanolamine (anandamide. AEA) and their metabolic enzymes (Devane et al., 1992; Di Marzo et al., 1999; Mechoulam et al., 1995; Munro et al., 1993) – is a major modulator of neuronal plasticity in the central nervous system(Freund et al., 2003). The eCB system has recently emerged as a potential target in the therapy of PTSD (Hill et al., 2018) as i.) its components are abundantly expressed within the neuronal circuits relevant to fear learning and anxiety, such as the medial prefrontal cortex (mPFC), amygdala (AMY) and hippocampus (HC) (Herkenham et al., 1990; Matsuda et al., 1990); ii.) it substantially contributes to the regulation of cognitive and emotional functions (Zanettini et al., 2011), both significantly altered in PTSD; iii.) its neuropharmacological manipulation in rodent models of PTSD have been shown to affect the behavioral consequences of trauma exposure (Aliczki and Haller, 2015; Danandeh et al., 2018; Fotio et al., 2023); and iv.) reports on PTSD patients self-medicating with cannabis suggest that symptoms can be ameliorated through activation of the eCB system (McNabb et al., 2023).

While several studies aimed to clarify the involvement of eCB function in different PTSD symptoms by employing means of neuropharmacological approaches in laboratory rodents, our knowledge on the specific trauma-induced changes in the components of the eCB system and their direct contribution to the development of PTSD symptoms is relatively scarce and often contradicting. There is limited information available on the expression patterns of cannabinoid receptor type-1 (CB_1_R) in PTSD but the studies addressing the issue conclusively showed that expression is elevated in PTSD patients (Neumeister et al., 2015, 2013). There are significantly more studies on the levels of eCBs in PTSD, however, findings on this issue are inconclusive and very few studies addressed the potential link between eCB levels and PTSD susceptibility. Several reports investigating circulating plasma eCB levels in connection with PTSD showed that AEA levels are chronically decreased in human PTSD patients and animal models of trauma exposure when compared to healthy, non-traumatized controls (Hill et al., 2013; Morena et al., 2018; Neumeister et al., 2013), although higher levels (Gunduz-Cinar et al., 2013; Hauer et al., 2013; Olango et al., 2012) or no detectable changes (Schaefer et al., 2014) were also reported. AEA content of PTSD patients (i.e. the susceptible subpopulation), when compared to trauma-exposed healthy individuals (i.e. the resilient subpopulation) was showed to be lower (Neumeister et al., 2013) suggesting that AEA content might predict susceptibility to develop PTSD following trauma exposure. Furthermore, as AEA content negatively correlated with symptom severity in PTSD patients (Bluett et al., 2014; deRoon-Cassini et al., 2022; Hill et al., 2013) one might hypothesize that low AEA levels are potentially involved in the pathophysiology of PTSD. In the case of posttraumatic 2-AG levels, studies are rather inconclusive as elevated (Hauer et al., 2013; Lim et al., 2016; Olango et al., 2012), decreased (Hill et al., 2013) and unchanged levels (Neumeister et al., 2013; Schaefer et al., 2014) were all reported in PTSD patients or traumatized laboratory rodents compared to non-traumatized healthy controls. Regarding susceptibility, the link between 2-AG content and PTSD susceptibility seems similarly inconclusive, as both elevated (Hauer et al., 2013) and decreased (Hill et al., 2013) serum 2-AG levels were reported in PTSD patients compared to trauma-exposed healthy individuals. Interestingly. deRoon-Cassini and colleagues suggested that 2-AG is differentially involved in PTSD pathophysiology than AEA, as its circulating content is positively correlated with symptom severity (deRoon-Cassini et al., 2022). Taken together, while eCB content was shown to be altered in PTSD, the trauma-induced alterations seem to be discrepant, especially when susceptibility issues are not taken into account and PTSD patients are compared with non-traumatized controls. As eCB levels and susceptibility were shown to be linked by several studies and a potential causal connection was suggested by correlations of eCB levels and symptom strength, one might hypothesize that individual differences in eCB function can determine susceptibility to develop trauma-induced behavioral symptoms. The putative connection between eCB function and PTSD susceptibility was suggested by the study of Danan and colleagues on laboratory rodents as well, showing that abnormal eCB reactivity to trauma contributed to subsequent susceptibility to develop a PTSD-like phenotype (Danan et al., 2021).

In the present study, we employed a rat model of PTSD to assess the putative link between eCB function and PTSD susceptibility in specific regions of the fear circuitry. We focused on generalized fear responses as a core symptom that allows sufficient variability to identify resilient and susceptible subpopulations. Through this model, our aims were twofold: i) to evaluate disparities in eCB levels among these subgroups, and ii) to scrutinize alterations in gene expression related to the eCB system and their potential association with susceptibility to trauma-induced adverse behaviors. Additionally to eCB-system related genes, we examined iii.) expression of several immediate early genes, as markers of neuronal activity (Hoffman et al., 1993), and modulators of neuroplasticity (Minatohara et al., 2015). Expression of such genes directly contributes to fear learning (Ploski et al., 2011), are involved in stress susceptibility (Kozlovsky et al., 2008) and their expression changes are associated with stress-related mental disorders (Selçuk et al., 2023). Furthermore, they interact with eCB signaling in the modulation of synaptic activity (Hartzell et al., 2018). Finally, leveraging our gene expression data, we employed unsupervised and semi-supervised machine learning-based statistical analysis iv.) to identify patterns of gene expression changes with strongest predictive capacity concerning PTSD susceptibility.

## 2. Material and Methods

### 2.1. Subjects

Subjects were adult male Long-Evans rats obtained from a breeding colony at the Medical Gene Technology unit of the Institute of Experimental Medicine established from rats from Charles River Laboratories (Italy). Animals were kept in Eurostandard Type IV cages measuring 59.8 cm x 38 cm x 20 cm. in groups of 3-4 rats per cage. In order to avoid stress contagion (stressed subject affecting the behavior of non-stressed controls) and social buffering (non-stressed control subjects affecting the behavior of stressed subjects) all subjects within the same cage underwent identical treatment (either fear conditioning or exposure to the context without traumatization. see Behavioral Procedures for details). In the animal rooms, temperature was maintained at 22±1°C, and relative humidity was kept at 20±10%. Laboratory chow and water were provided *ad libitum*. Rats were kept on a reverse 12-hour light/dark cycle with lights turning on at 8 p.m. and turning off at 8 a.m. The experiments followed the guidelines of the European Communities Council Directive of November 24. 1986 (86/609/EEC) and were conducted under license PE/EA/141-2/2020 issued by the Pest County Government Office.

### 2.2. Behavioral procedures

All behavioral procedures were carried out in the first part of the dark phase in dedicated experimental rooms by experimenters blind to experimental groups. Behavioral procedures were video recorded with Sony FDR-AX33 digital camcorders for later analysis.

#### 2.2.1. Contextual fear conditioning

Subjects were transported to a brightly lit experimental room (300 lux) and placed in a transparent Plexiglas chamber (measuring 40 x 40 x 40 cm) with a stainless steel electric grid floor. Following a 2.5 minute habituation period, a series of 10 scrambled electric footshocks were administered through the grid floor, each lasting 2 seconds and with an amperage of 2.4 mA, and separated by 30 second long periods between each shock. The conditioning session lasted for a total of 7 minutes. Non-shocked control subjects were introduced to the test chamber for 7 minutes without footshock delivery. Chambers were cleaned with 20% ethanol and dried with paper towel after each subject.

#### 2.2.2. Testing for contextual conditioned fear

28 days after footshock delivery. subjects were exposed to a chamber identical to the one used for conditioning (*Context A*) under the same experimental conditions without further footshock delivery for 5 minutes to assess contextual conditioned fear responses. The 28th day as the time point when behavior was first assessed was selected in order to model the lasting trauma-induced adverse behavioral changes in PTSD(American Psychiatric Association, 2022).

#### 2.2.3. Testing for generalization of contextual fear

29 days after footshock delivery (24 hours after *Context A* exposure), subjects were placed in a novel neutral context (*Context B*) for 20 minutes. *Context B* was carefully designed to differ entirely from *Context A*. utilizing different visual, olfactory, and tactile stimuli: i.) the experimenter was different from the one conducting conditioning and *Context A* exposure. ii.) a different experimental room was used, illuminated with dim red light (5 lux). iii.) plastic inserts were placed over the grid floor and the walls to modify shape, floor and wall color of the chambers. iv.) chambers were cleaned between subjects using soapy water.

#### 2.2.4. Behavioral analysis

Video recordings taken during behavioral procedures were analyzed using the Noldus Ethovision XT15 (Noldus Information Technology. The Netherlands) software, in which the activity monitoring feature was used to quantify fear responses by calculating the percentage of time spent freezing, a typical fear response in rodents (Fanselow, 1980). This feature is based on an automated, frame-by-frame analysis of behavior video recordings during which the algorithm compared the pixel differences between subsequent frames. Pixel differences below a specific *a priori* set threshold represented the immobility of the subject (i.e., freezing behavior). Thresholds were constant across subjects within an experiment and were compared to and validated by previous manual behavior analyses of randomly selected subjects conducted by trained experimenters blind to treatment groups.

### 2.3. Identification of resilient and susceptible subpopulations

Generalized fear responses in a neutral context measured by the percentage of time spent freezing during the *Context B* test were used to identify resilient and susceptible individuals. Considering the estimated proportion of patients diagnosed with PTSD and healthy trauma-exposed individuals in the populations (Breslau et al., 1998, 1991), we characterized a resilient and susceptible subpopulation as the upper and lower quartile of the population based on their fear generalization. This approach allowed for a clear distinction between the two groups and aided in further analysis of their behavior and physiological functions. Animals that fall between susceptible and resilient categories were classified as intermediate individuals.

### 2.4. High-performance liquid chromatography-tandem mass spectrometry (HPLC/MS/MS)

Subjects were rapidly decapitated, and their brain was manually removed. Brains were cooled on wet ice and were dissected into 2 mm thick coronal slices on a stainless steel brain mold. Tissue blocks containing relevant areas in the neurobiological basis of PTSD, the prelimbic (PrL) and infralimbic (IL) cortices of the mPFC, basolateral amygdala (BLA). ventral hippocampus (vHC) were identified according to the atlas of Paxinos and Watson (Paxinos and Watson, 2006) based on clearly visible anatomical reference points (e.g. forceps minor and piriform cortex for the mPFC; the internal and external capsule as well as the optic tract, and the commissure of the stria terminalis for the BLA). In addition to these landmarks, subtle color patterns that circumscribed particular brain regions also guided dissection. The sampled regions were dissected bilaterally and pooled within each subject. Samples were moved into sterile Eppendorf tubes and snap-frozen on dry ice. Tissue samples were kept at - 80°C until later analysis. Sample weight was calculated from the difference in tube weight measured before and after sampling using an analytical balance (Kern ABJ 80-4NM. Kern-Sohn GmbH., Germany). Sample preparation and eCB measurement was conducted as described earlier by Lee and colleagues (Lee et al., 2015). First, tissue samples were incubated on ice for 30 min in 1 ml of HPLC-gradient grade methanol (Merck KGaA, Darmstadt, Germany) containing the deuterated internal standards 2-arachidonoylglycerol-d5 (100 ng/ml) and arachidonoylethanolamide-d (1 ng/ml) (Cayman Chemical, Ann Arbor, MI, USA). Samples were homogenized and centrifuged on an Eppendorf miniSpin microtube centrifuge at 13,400 rpm for 15 min. The supernatant was transferred into a plastic tube and was diluted with 3ml of ultrapure water. The samples were subjected to solid-phase extraction (SPE) sample clean-up according to the following protocol. First, the SPE cartridges (SUPELCO Discovery DSC-18 1 ml tubes, 100 mg) were conditioned with 2 ml of methanol and 2 ml of ultrapure water. Diluted samples were loaded at approximately 0.5 ml/min flow rate to apply a gentle vacuum. In the consecutive washing step, 2 ml of ultrapure water and 2 ml of methanol/water (50:50 v/v%) were percolated through the sorbent material. Elution was performed with 0.7 ml of methanol. Eluates were diluted to initial HPLC eluent composition with 10 mM ammonium formate (Honeywell Riedel-de Haen GmbH, Seelze, Germany) solution. To measure eCB levels, a PerkinElmer Life and Analytical Sciences HPLC Series 200 system was employed coupled to an Applied Biosystems/Sciex 4000 QTRAP triple quadrupole/linear ion trap tandem mass spectrometer operated in positive electrospray ionization mode. The electrospray ionization ion source parameters were set as follows: curtain gas, 10; ion spray voltage, 5000 V; temperature, 500°C; collisionally activated dissociation gas, medium; gas 1, 50; gas 2, 40. Chromatographic separation was achieved with a Phenomenex Kinetex C18 column (50mm x 3.00 mm, 2.6 x m, 100 Å) using methanol and 10 mM ammonium formate as elution solvents at a flow rate of 500 µl/min and injection volume of 50 µl. The initial eluent condition was 80% methanol/20% buffer that was increased to 85% organic phase during 3 min and then further elevated to 95% during 2 min and was kept at this condition for 2 min. Afterward, the column was equilibrated to the initial condition. Analytes were detected in multiple reaction monitoring (MRM) mode at the following ion transitions and parameter settings: (1)2-AG, MRM transition [mass/charge ratio (m/z), 379.4 287.2, 379.4 91.1], declustering potential (81 V), collision energy (23 V, 81 V), collision cell exit potential (10 V, 8 V); (2) 2-arachidonoylglycerol-d5, MRM transition (m/z, 384.4 287.2, 384.4 91.1), declustering potential (81 V), collision energy (23 V, 81 V), collision cell exit potential (10 V, 8 V); (3) arachidonoylethanolamide, MRM transition (m/z, 348.4 62.1, 348.4 90.9), declustering potential (51 V), collision energy (43 V, 63 V), collision cell exit potential (4 V, 8 V); and (4) arachidonoylethanolamide-d4, MRM transition (m/z, 352.4 66.0, 352.4 91.2), declustering potential (81 V), collision energy (41 V, 77 V), collision cell exit potential (6 V). The peak areas were determined with Analyst 1.4.2. software. The quantity of the analytes was calculated by comparing their peak areas with those of the deuterated internal standards, and normalized to sample weight.

### 2.5. Real-Time quantitative PCR (RT-qPCR)

Rats were rapidly decapitated, and the brains were promptly excised from the skull, cooled using ice, and maintained in an RNase-free environment. Based on the atlas of Paxinos and Watson (Paxinos and Watson, 2006), a 2 mm thick coronal section containing the respective mPFC and the AMY region was sliced on a stainless steel mold. Whole HC samples were collected using ophthalmic tweezers. The mPFC, AMY, and HC samples were dissected promptly, placed in sterile Eppendorf tubes, and snap-frozen on dry ice. The tissue samples were stored at -80°C until later analysis.

The samples were moved into QIAzol lysis reagent (Qiagen. The Netherlands) and homogenized using a Tissue-Tearor (BioSpec Products. USA). The total RNA extraction was carried out using the RNeasy Lipid Tissue Mini Kit (Qiagen. The Netherlands) according to the manufacturer’s protocol. All samples were treated with RNase Free DNase Set (Qiagen. The Netherlands) to remove genomic DNA contamination. The Qubit 4 Fluorometer (Invitrogen. USA) was used to obtain information about the quantity (Qubit RNA BR Assay Kit; Invitrogen. USA) and quality (Qubit RNA IQ Assay Kit; Invitrogen. USA) of the RNA in the samples. The total RNA samples were diluted in nuclease-free water to a concentration of 0.1 µg/µl. using 1 µg of RNA to perform reverse transcription.

Each sample underwent reverse transcription of 1 µg of total RNA using random primers and the High Capacity cDNA reverse transcription kit (Thermo Fisher Scientific. USA). Reverse transcription was performed on a VWR Ristretto thermal cycler (VWR International GmbH. Germany). The cDNA concentrations were measured using a Qubit ssDNA assay kit (Invitrogen. USA) on a Qubit 4 Fluorometer.

The cDNA samples were diluted in nuclease-free water and then combined with TaqMan® Gene Expression Master Mix to create a PCR reaction mix, resulting in a final concentration of 1 ng/µl of cDNA. The expression of 45 genes associated with either the eCB system or neuronal activity and 3 housekeeping genes were analyzed using Custom 384-well microfluidic TaqMan® Gene Expression Array Cards (Applied Biosystems. USA). The array cards consisted of eight fill reservoirs linked to a series of 48 reaction wells. These wells contained dried-down TaqMan® assays for 48 unique genes. PCR reaction mixes were loaded and centrifuged twice at 331 x g for 1 minute. The main distribution channels of the array card were then sealed with a TaqMan® Array Card Sealer to preserve the mixtures. A ViiA™ 7 Real-Time PCR System (Applied Biosystems. USA) was used to perform RT-qPCR. The cycle threshold (Ct) values and other data related to PCR reactions were collected using the QuantStudio software (Applied Biosystems. USA). The gene expression levels for the candidate genes were normalized to the geometric mean of expression levels of the three endogenous control genes (*18S*, *Actb*, and *Gapdh*). The relative quantity was then calculated using the 2^-ΔΔCt^ method (Livak and Schmittgen, 2001).

### 2.6. Experimental design

In *Experiment 1*, we evaluated the variations in AEA and 2-AG levels over time after a traumatic experience by assessing the short- (3 hours after footshocks) and long-term (28 days after footshock) effects of shock exposure. Subjects were assigned to experimental groups *home cage control*, *non-shocked control,* and *shocked*. Following footshock delivery, shocked and non-shocked control subjects were returned to their home cages and sacrificed either 3 hours or 28 days later under baseline conditions. Home cage control subjects not exposed to the footshock apparatus were sacrificed at each time point as well. Brain punch samples from PrL, IL, vHC, and BLA were collected for eCB content assessment by HPLC/MS/MS.

In *Experiment 2*, we assessed the long-term effects of traumatic experiences on AEA and 2-AG levels in connection with resilience/susceptibility to develop trauma-induced generalized fear responses, a core symptom of PTSD. We subjected rats to contextual fear conditioning, and then 28 days later, a contextual reminder (*Context A*) was performed to assess conditioned fear responses. On the 29th day post-conditioning, subjects were exposed to a different context (*Context B*) to evaluate fear generalization. After *Context B* exposure, subjects were immediately sacrificed, and brain punch samples from PrL, IL, vHC, and BLA were collected. AEA and 2-AG levels were assessed by HPLC/MS/MS in resilient and susceptible subpopulations that were identified *a posteriori* based on generalized fear responses in *Context B*. Additionally, a cohort of non-shocked control subjects underwent exposure to Contexts A and B without the presentation of footshocks on the initial day of conditioning.

In *Experiment 3*, we assessed trauma-induced expression patterns in eCB system-associated genes in connection with resilience/susceptibility to develop trauma-induced generalized fear responses. Following contextual fear conditioning and subsequent exposure to *Context A* and *Context B*, subjects were classified into susceptible and resilient groups, as outlined in *Experiment 2*. Rats were sacrificed immediately following *Context B* exposure, and brain punch samples were collected from the mPFC, HC, and AMY regions. Punch samples were used for RT-qPCR analysis of the expression of eCB system-related genes.

### 2.7. Statistical analysis

#### 2.7.1. Descriptive statistics and hypothesis testing

Behavioral and eCB level data are shown as mean ± standard error of the mean. eCB content in *Experiment 1* was evaluated by one-factor analysis of variance (ANOVA) (factor: *experimental group*). Time spent with freezing during conditioning and contextual reminders as well as brain eCB content in *Experiment 2* were evaluated by one-factor ANOVA (factor: *experimental group*). Tukey’s multiple comparisons tests were performed for post-hoc analyses when a main effect was significant. Correlations between brain eCB levels and time spent freezing during *Context B* exposure were evaluated by the Pearson correlation coefficient. For the analyses including all individuals in *Experiment 2*, the Mann-Whitney U tests were performed. For the analyses of PCR data in *Experiment 3*, we used multiple unpaired t-tests. P-values lower than 0.05 were considered statistically significant. All statistical analyses above were conducted with GraphPad Prism software version 9.5.1 (MA. USA).

#### 2.7.2. Principal component analysis

To assess the structure and correlatedness of the expression data of different brain regions, we conducted a principal component analysis (PCA) using the varcomp R package. We calculated the proportion of the variance explained by each principle component to see the dimensionality of the expression data. We also calculated the loadings of gene expression variables on each principle component to reveal their correlatedness.

#### 2.7.3. Random Forest classification

To decrease the dimensions of our multivariate data and identify the gene expression predictors of resilient and susceptible subpopulations, we applied the semi-supervised machine learning method. Random Forest (RF) (Breiman, 2001). RF ranks the importance of variables in the classification of subjects to subpopulations, in this particular case, to resilient and susceptible subgroups. It can create thousands of decision trees and splits subjects into groups according to the expression of a random subset of the genes. This way RF is suitable for classifying samples that have many features, even with limited sample sizes. Here. i) we developed a saturated model using all gene expression data of all investigated brain subregions and ii) ranked the genes according to their importance. Then we iii) created subset models with the most important variables, one in which all areas were included and three in a subregion-specific manner. Finally. iv) we compared these subset models according to their classification accuracy. Note that, due to the outstanding prediction accuracy of the gene *Npas4* expression in the HC and that RF can be biased by features with the impact of this level, we conducted two separate analyses, one with and one without the inclusion of HC *Npas4* data. All models produced 5.000 decision trees and considered 9 genes at a time. Sampling was balanced to 7-7 subjects per group in the RF analysis. Variable importance was ranked based on permutation importance, which shows the mean decrease in prediction accuracy of a feature in response to randomly permuting its values.

## 3. Results

### 3.1. Basal eCB content does not reflect previous trauma exposure in the whole population

To study the short- and long-term effects of trauma exposure on basal levels of 2-AG and AEA in brain regions involved in the regulation of fear responses and memory, we obtained brain samples at either 3 hours or 28 days following fear conditioning (Figure 1A). No significant differences in eCB concentration were observed between groups at the studied time points and brain areas. These findings suggest that basal eCB levels in a particular brain subregion do not reflect previous exposure to a traumatic event at the studied time points when assessed in the whole population (Figure 1B-E). For data of statistical analyses, see Table 1.

**Figure 1.**
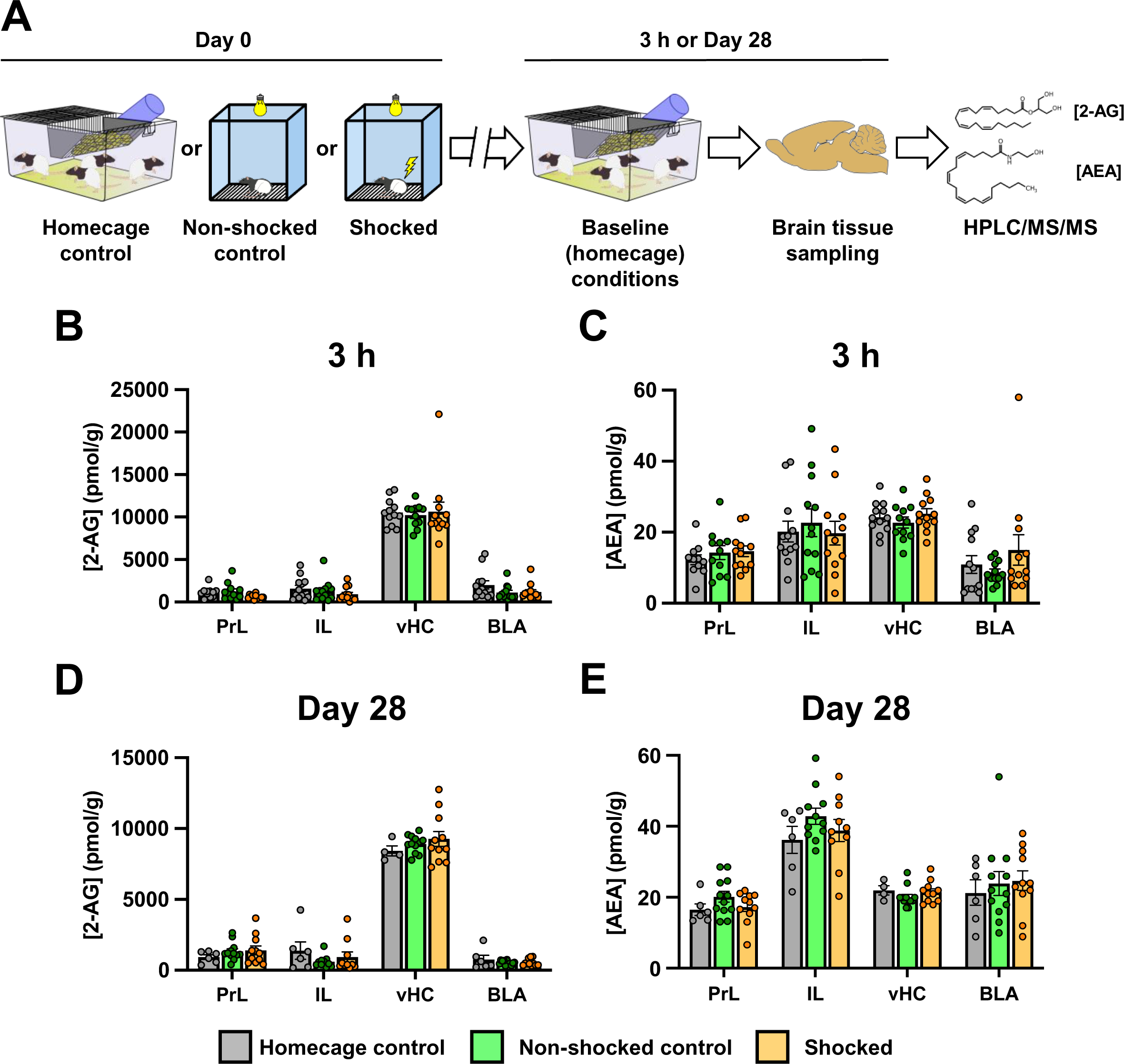
Basal endocannabinoid content does not reflect previous trauma exposure in the whole population. **A**) Experimental design schematic showing experimental groups: control subjects left undisturbed in their homecage without undergoing behavioral testing, non-shocked control subjects exposed to the footshock context without delivery of footshocks, and subjects exposed to footshocks. Subjects were sacrificed under baseline conditions from their home cage at either 3 h or 28 days after footshock/context exposure. Brain samples from prelimbic cortex (PrL), infralimbic cortex (IL), ventral hippocampus (vHC) and basolateral amygdala (BLA) were collected for HPLC/MS/MS analysis of 2-arachidonoylglycerol (2-AG) and N-arachidonoylethanolamin (AEA) levels. **B-E**) Neither footshock delivery, nor context exposure led to changes in 2-AG or AEA levels in the assessed regions. Data are presented as mean ± SEM. One-way ANOVA.

**Table 1.** Statistical data of *Experiment 1*. 2-AG: 2-arachidonoylglycerol; AEA: N-arachidonoylethanolamine; PrL: prelimbic cortex; IL: infralimbic cortex; vHC: ventral hippocampus; BLA: basolateral amygdala; ANOVA.

| sampling time point | area | 2-AG |  | AEA |  |
| --- | --- | --- | --- | --- | --- |
|  |  | F(DFn, DFd) | p-value | F(DFn, DFd) | p-value |
| 3h | PrL | F(2, 31) = 2.972 | 0.066 | F(2, 31) = 0.604 | 0.552 |
|  | IL | F(2, 31) = 1.056 | 0.359 | F(2, 33) = 0.215 | 0.807 |
|  | BLA | F(2, 32) = 0.0677 | 0.934 | F(2, 32) = 1.154 | 0.328 |
|  | vHC | F(2, 33) = 1.933 | 1.933 | F(2, 33) = 1.187 | 0.317 |
| 28 days | PrL | F(2, 33) = 1.187 | 0.461 | F(2, 26) = 1.569 | 0.227 |
|  | IL | F(2, 24) = 1.176 | 0.325 | F(2, 24) = 1.241 | 0.307 |
|  | BLA | F(2, 24) = 0.710 | 0.501 | F(2, 24) = 0.964 | 0.395 |
|  | vHC | F(2, 26) = 0.810 | 0.455 | F(2, 26) = 0.222 | 0.801 |

### 3.2. Brain subregion-specific eCB content is linked to susceptibility to developing trauma-induced symptoms

In our second experiment, we evaluated the levels of 2-AG and AEA in resilient and susceptible subpopulations along with non-shocked control subjects, following the testing of fear generalization (Figure 2A). Resilient and susceptible subpopulations were classified based on the duration spent with freezing behavior in the neutral context (*Context B,* i.e., generalized fear). Subjects in the upper quartile were identified as susceptible, while those in the lower quartile were categorized resilient. During *Context B*. we observed significant differences in freezing duration between these groups. Post-hoc comparisons revealed that susceptible individuals froze significantly longer than the resilient and non-shocked groups. In contrast, resilient subjects exhibited no difference in freezing duration compared to non-shocked control subjects (Figure 2D). We found no difference in freezing duration between resilient and susceptible subjects during contextual fear conditioning (Figure 2B) or contextual reminder (*Context A,* Figure 2C). However, both subpopulations exhibited increased freezing duration compared to non-shocked controls. When we compared all shocked animals (including the intermediate quartiles) to non-shocked controls, we observed a significant increase in freezing duration during both contextual fear conditioning and contextual reminder (*Context A,* Supplemental Figure 1A and 1B), but interestingly, we detected no significant difference in freezing duration in *Context B* (Supplemental Figure 1C). This lack of difference is possibly due to high variability in generalized fear among subjects previously exposed to footshocks.

**Figure 2.**
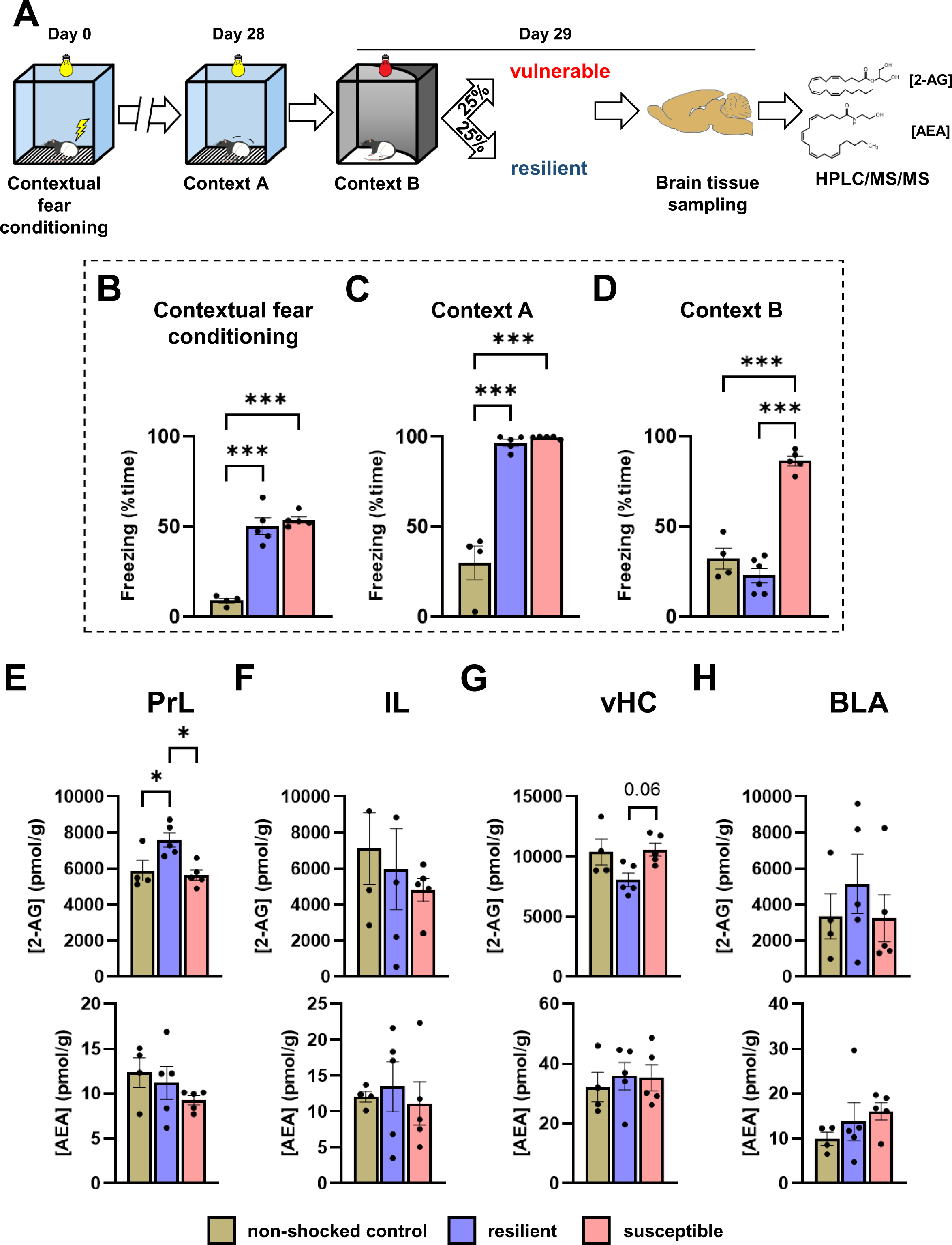
Brain subregion-specific endocannabinoid content is linked to susceptibility to developing trauma-induced symptoms. **A**) Experimental design schematic showing our model of trauma exposure. Following contextual fear conditioning (Day 0), contextual fear memory expression was assessed 28 days later in the traumatic context (*Context A*. Day 28). The next day, fear generalization was studied in a novel, safe context (*Context B*. Day 29). Resilient and susceptible subjects were categorized based on the upper and lower quartiles of generalized fear characterized by freezing duration in *Context B*. Subjects were immediately sacrificed after *Context B* testing, and brain samples from prelimbic cortex (PrL), infralimbic cortex (IL), ventral hippocampus (vHC) and basolateral amygdala (BLA) were collected for HPLC/MS/MS analysis of 2-arachidonoylglycerol (2-AG) and N-arachidonoylethanolamin (AEA) levels. Non-shocked controls underwent the same procedure without the delivery of footshocks. **B**) Contextual fear conditioning induced freezing behavior during the conditioning session. (**C**) which remained high during *Context A* contextual reminder in resilient and susceptible groups compared to the non-shocked control. **D**) The susceptible group showed markedly high fear generalization, indicated by the higher freezing responses compared to resilient or non-shocked control groups. **E**) The resilient group exhibited higher levels of 2-AG in the PrL than susceptible and non-shocked control groups, without changes in AEA levels. **F**) 2-AG and AEA levels did not show differences in the IL. **G**) In the vHC, there was a significant main effect in the 2-AG levels without significant differences in pairwise comparisons, although 2-AG showed a tendency to decrease in resilient subjects. AEA levels did not differ between groups in this region. **H**) 2-AG and AEA levels in the BLA did not show differences. *: p < 0.05. **: p < 0.01. ***: p < 0.001; Data are presented as mean ± SEM. One-way ANOVA.

Our analysis of eCB content in specific brain regions demonstrated a significant effect of the experimental group on 2-AG levels in the PrL. Post-hoc analysis revealed that resilient subjects exhibited significantly higher 2-AG levels compared to the non-shocked and susceptible groups. However, we found no differences in AEA content in this region (Figure 2E). In IL, we observed no significant group differences in 2-AG or AEA content (Figure 2F). In the ventral hippocampus (vHC), there was a strong trend towards reduced 2-AG levels in resilient animals (Figure 2G), but AEA levels showed no differences in this brain region (Figure 2G). For data of statistical analyses see Table 2. We found no group differences in the BLA regarding 2-AG or AEA levels (Figure 2H).

**Table 2.**
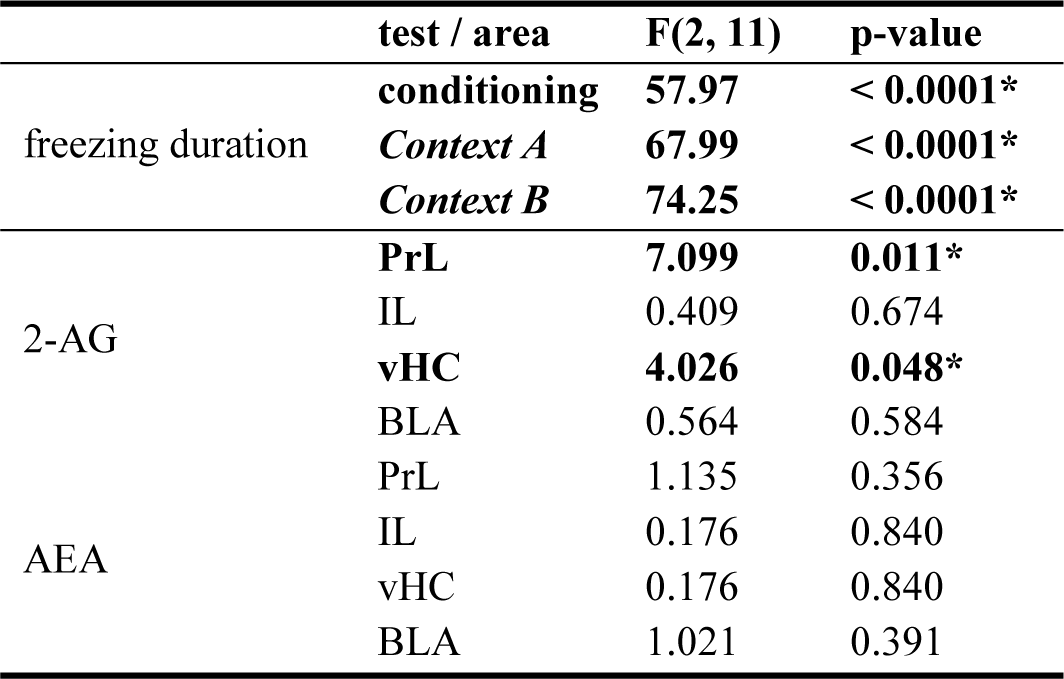
Statistical data of *Experiment 2*. 2-AG: 2-arachidonoylglycerol; AEA: N-arachidonoylethanolamine; PrL: prelimbic cortex; IL: infralimbic cortex; vHC: ventral hippocampus; BLA: basolateral amygdala; *: significant group effect. ANOVA.

We also investigated variations in 2-AG or AEA levels by comparing all subjects who received footshocks (with all shocked subjects pooled together) to non-shocked controls. This investigation did not reveal significant differences at the studied brain sites (Supplemental Figure 1E-H, for data on statistical analyses, see Supplemental Table 1).

As we showed that eCB content reflects susceptibility in our model, we assessed whether populational variability of eCB levels correlate with the variability of generalized fear. We observed a significant positive correlation between total freezing duration and 2-AG concentration in the vHC of subjects who received footshocks (i.e. the whole sample, including ‘intermediate’ animals). (Figure 3. for data of statistical analyses. see Table 3).

**Figure 3.**
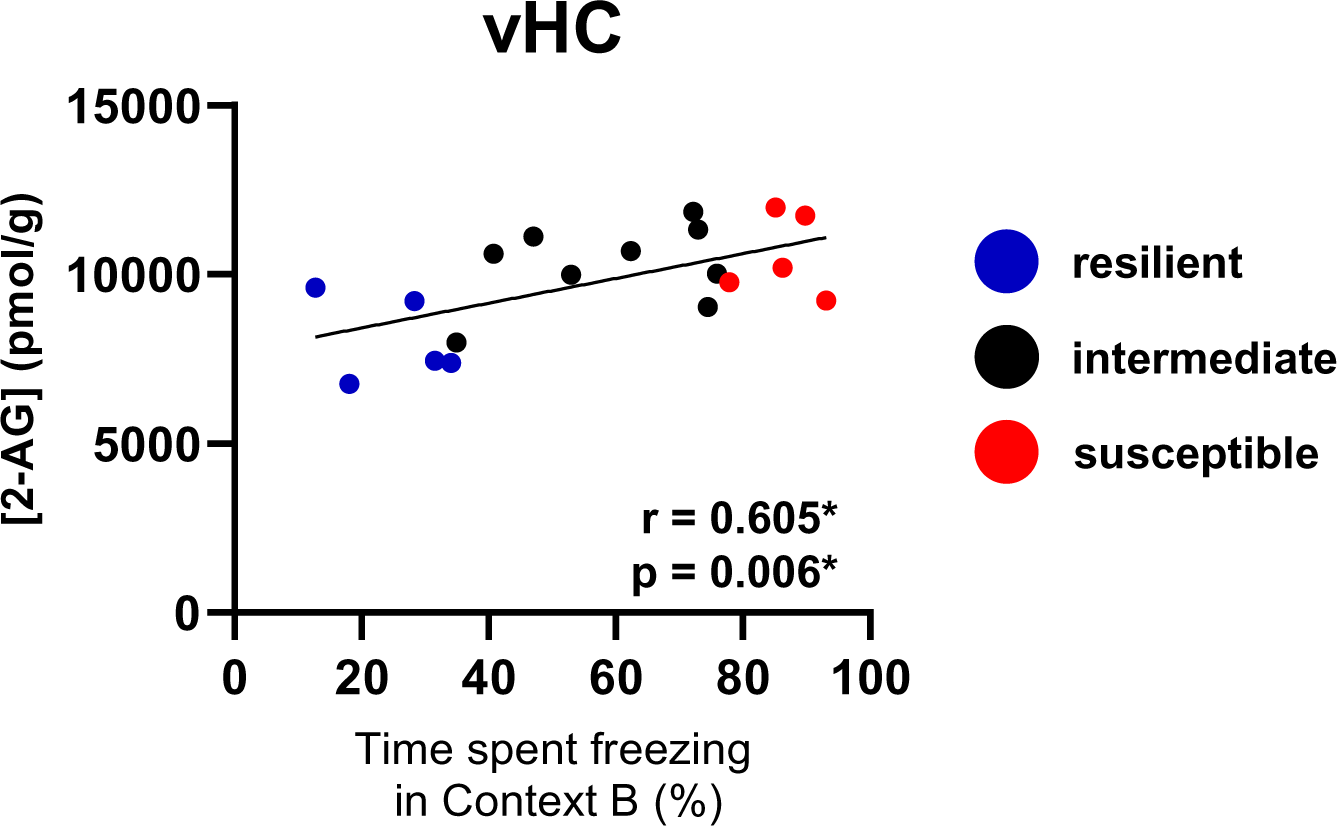
2-arachidonoylglycerol (2-AG) levels in the ventral hippocampus correlate with the strength of footshock-induced generalized fear responses. In the ventral hippocampus (vHC), there was a significant positive correlation between 2-AG levels and the time spent freezing in *Context B*. Pearson correlation test.

**Table 3.** Statistical data of correlation analyses between generalized fear and endocannabinoid content in *Experiment 2*. 2-AG: 2-arachidonoylglycerol; AEA: N-arachidonoylethanolamine; PrL: prelimbic cortex; IL: infralimbic cortex; vHC: ventral hippocampus; BLA: basolateral amygdala; *: significant correlation. Pearson correlation coefficients.

|  | area | Pearson r | p-value |
| --- | --- | --- | --- |
| freezing duration in context B vs. 2-AG cc. | PrL | -0.165 | 0.498 |
|  | IL | -0.128 | 0.601 |
|  | **vHC** | **0.605** | **0.006\*** |
|  | BLA | -0.304 | 0.205 |
| freezing duration in context B vs. AEA cc. | PrL | -0.301 | 0.210 |
|  | IL | -0.098 | 0.689 |
|  | vHC | -0.195 | 0.423 |
|  | BLA | -0.164 | 0.500 |

### 3.3. Susceptibility to generalize learned fear is associated with fear circuitry neural hypoactivity and altered neuroplasticity- and eCB system-related gene expression

In our subsequent experiment, we employed RT-qPCR analysis to evaluate the gene expression patterns related to the eCB system in key areas of regulating fear responses in subjects classified as resilient and susceptible. We targeted 40 genes essential to the eCB system, encompassing receptors, enzymes implicated in eCB synthesis and metabolism, transporters, and proteins involved in eCB signaling pathways. Additionally, we examined the expression of 5 markers of neuronal activation and neuroplasticity to assess the neuronal activity patterns within the circuitry (Figure 5A). The complete list of the 45 evaluated genes is provided in Supplemental Table 2.

Considering the extensive number of parameters from various brain areas that were assessed to elucidate the resilient and susceptible phenotype, we employed Principal Component Analysis (PCA) for dimension reduction. We found that the first three components explained 57% of the total variance, while each subsequent component accounted for less than 10% (Figure 4A). Therefore, we concentrated on these primary dimensions. The variable loadings of the first three components were distributed in a highly area-specific manner. Specifically, hippocampal genes positively loaded on the first component, amygdalar genes negatively loaded on the second component, and prefrontal genes negatively loaded on the third component of the total variability (Figure 4B). Given that gene expression data from different areas appeared uncorrelated, we implemented additional approaches to further reduce the dimensionality of our data and identify predictors of the resilient and susceptible phenotype. We applied the machine-learning-based Random Forest (RF) classification to categorize subjects into resilient and susceptible subpopulations. This allowed us to determine which gene products are the most potent predictors of such a phenomenon. HC *Npas4* expression emerged as the strongest predictor of the phenotype, exhibiting more than four-fold greater variable importance compared to the second most important parameter (Figure 4C top). To address potential bias, we conducted a supplementary analysis excluding *Npas4* (AMY) from consideration, revealing additional potential predictors: *Abhd12* (mPFC), *Daglb* (HC), *Fos* (HC), *Fosb* (HC), *Arc* (mPFC), *Faah* (mPFC), and *Slc17a8* (AMY) (refer to Figure 4C bottom). Utilizing these variables, we constructed a model incorporating all subregions, as well as specific models for individual subregions. Upon comparison using receiver operating curve analysis, gene expression analysis in the amygdala, prefrontal and hippocampal demonstrated varying levels of predictive power, with the hippocampal model exhibiting superior performance (Figure 4D). Hypothesis testing methods validated most of these predictors, as our analyses revealed a substantial downregulation of markers of neuronal activity and neuroplasticity associated with altered eCB-related gene expressions in susceptible compared to resilient subjects across the studied regions (Figure 5B-D). For detailed statistical data, please refer to Table 4.

**Figure 4.**
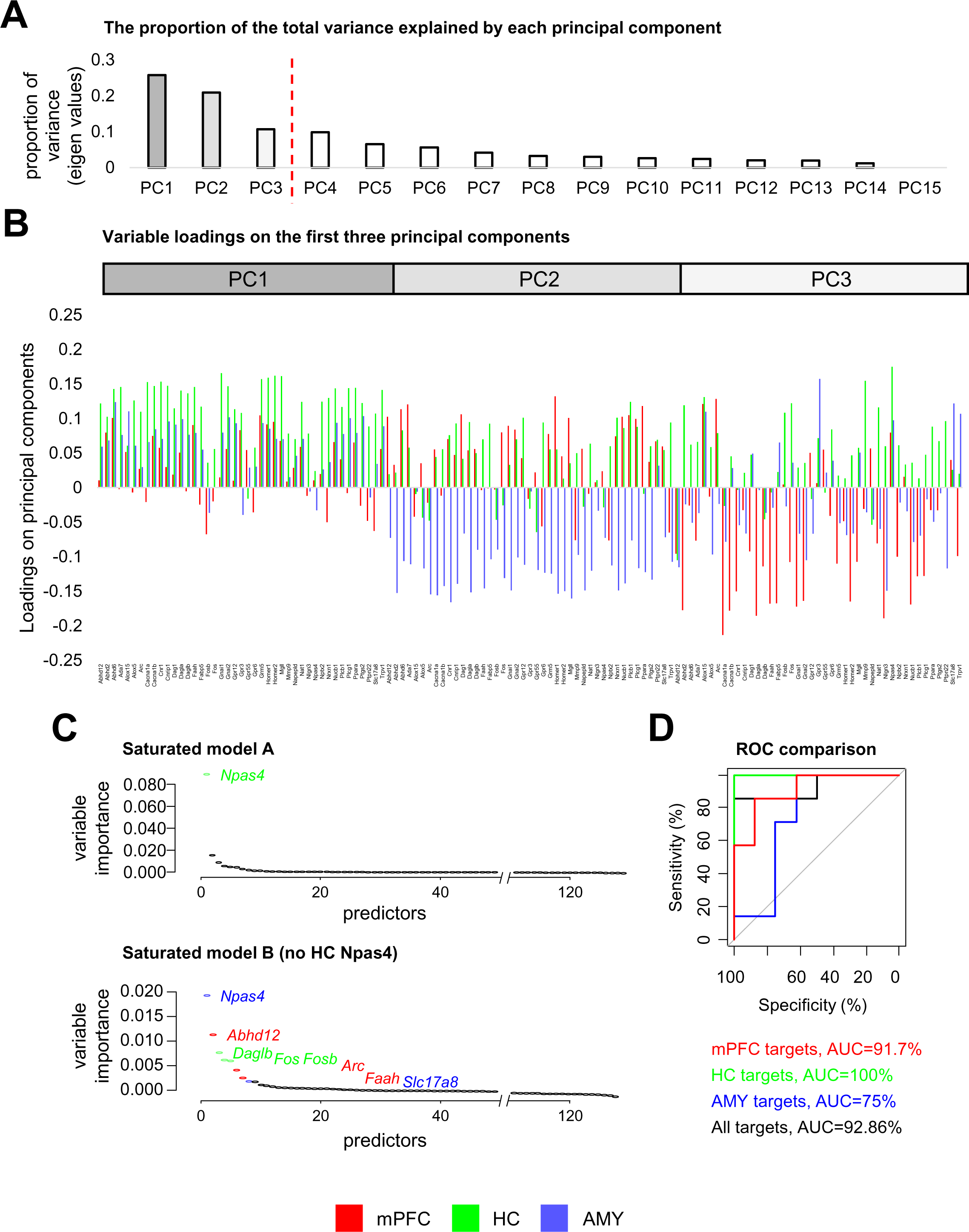
Expression of neuronal activity gene markers, neuroplasticity and endocannabinoid system-related genes predict susceptibility to generalize learned fear. **A**) Scree plot of the principal component analysis (PCA) involving all assessed 40 gene expression data of all 4 investigated brain areas. Eigen values show the proportion of variance of generalized fear in *Context B* explained by each component. Red dashed line indicates the last component that explains more than 10% of the total variability. **B**) Variable loadings on the main components (PC1-3, principal components 1-3). Variable-component correlations show an area-specific expression pattern. C) Variable importance levels in two saturated Random Forest models with (top, model A) and without (bottom, model B) the hippocampal *Npas4* expression data. D) ROC curves and the area under the curve (AUC) values of the reduced models involving the significant predictors of the B model of all (black) or in a given brain subregion (red: medial prefrontal cortex (mPFC); green: hippocampus (HC); blue: amygdala (AMY).

**Figure 5.**
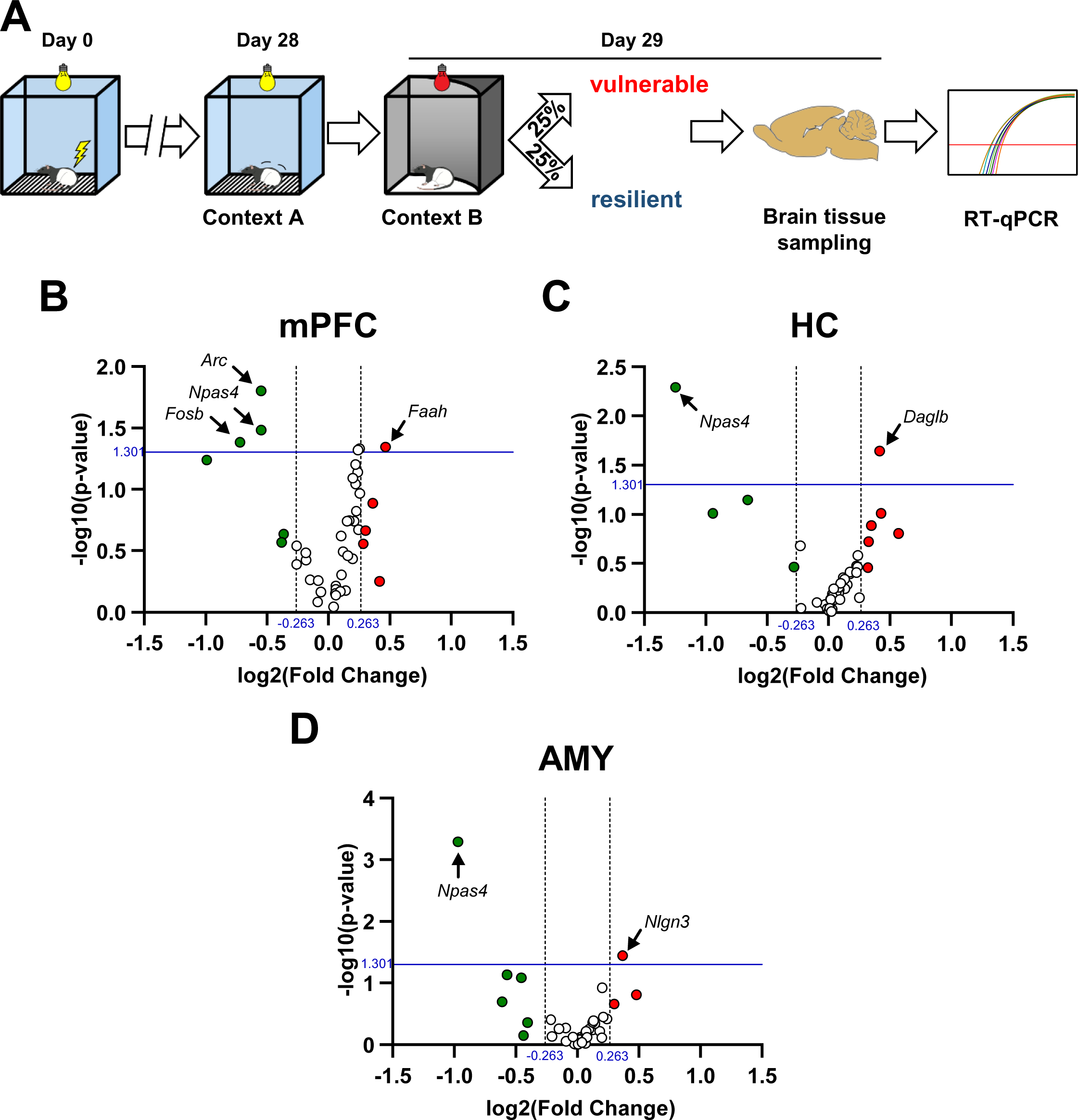
Susceptibility to generalize learned fear is associated with fear circuitry neural hypoactivity and altered neuroplasticity- and endocannabinoid system-related gene expression. **A**) Experimental design schematic showing our model of trauma exposure. Following contextual fear conditioning (Day 0), expression of contextual fear memory was assessed 28 days later in the traumatic context (*Context A*, Day 28). The next day, fear generalization was studied in a novel, safe context (*Context B*, Day 29). Resilient and susceptible subjects were categorized based on the upper and lower quartiles of the generalized fear spectrum characterized by freezing duration in *Context B*. Subjects were immediately sacrificed after *Context B* testing, and brain samples from the medial prefrontal cortex (mPFC), hippocampus (HC), and amygdala (AMY) were collected to analyze endocannabinoid (eCB) system-related gene expression using RT-qPCR. **B-D**) Gene expression differences and statistical significance levels of the susceptible group compared to the resilient group displayed by volcano plots in the mPFC (**B**), HC (**C**), and AMY (**D**). The horizontal blue lines represent the criteria of statistical significance p = 0.05. Genes that exhibit a fold change greater than 1.2 are denoted by green dots for downregulation and red dots for upregulation. Vertical dashed lines represent the margin of 1.2 fold change.

**Table 4.**
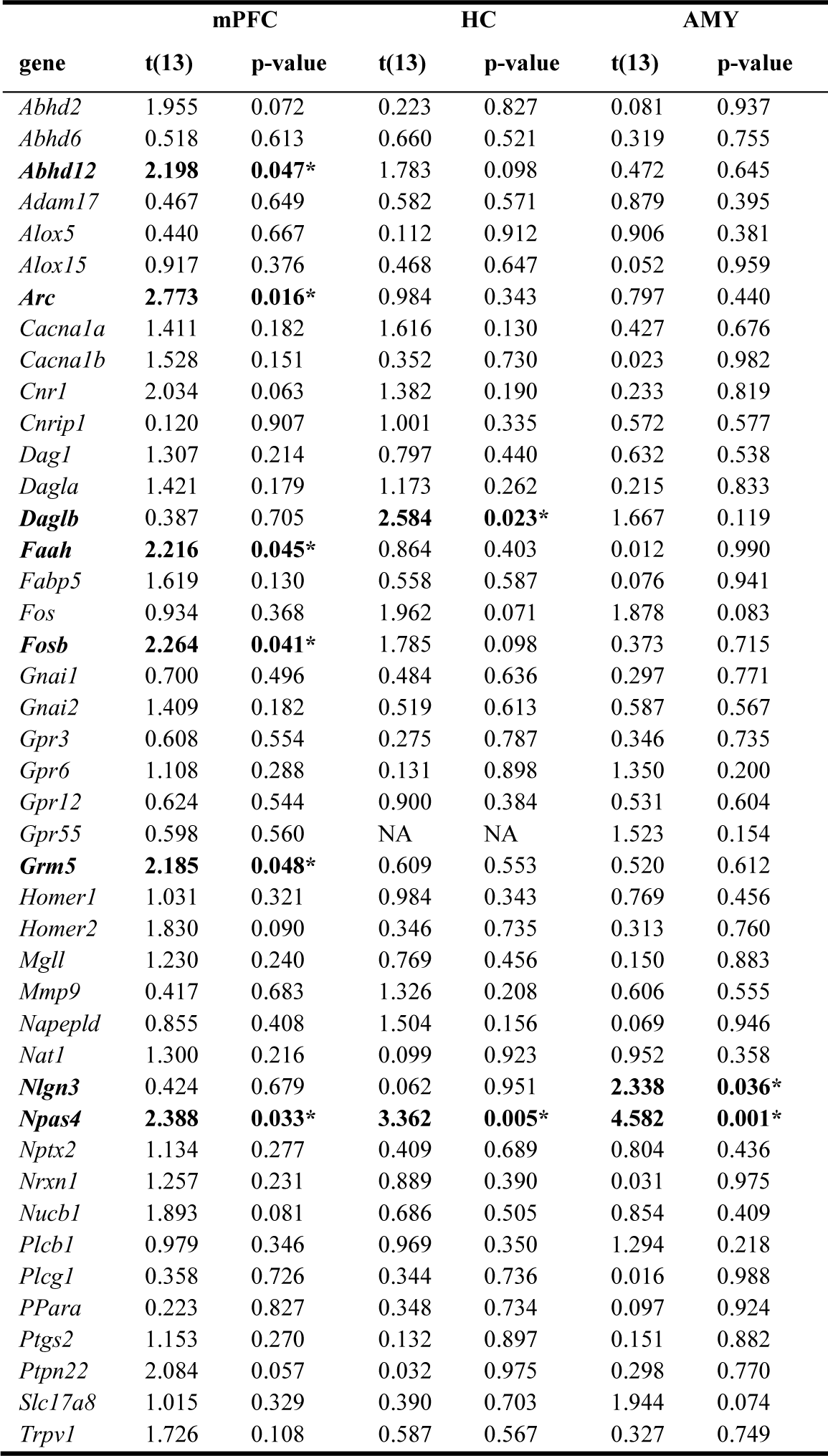
Statistical data of gene expression analysis in *Experiment 3*. *Cnr2* showed consistently low expression at all assessed regions, therefore this gene was excluded from analysis. mPFC: medial prefrontal; HC: hippocampus; AMY: amygdala; *: significant group effect. Multiple t-test.

## 4. Discussion

Our findings showed that basal 2-AG and AEA levels in the fear circuitry do not inherently reflect previous traumatic experiences when examined across the entire population. However, upon individual classification into susceptible or resilient groups regarding the development of trauma-induced fear generalization, distinct trauma-related disparities in eCB content within the PrL and vHC emerged. Specifically, susceptibility was associated with higher 2-AG content in the PrL and lower in the vHC. Notably, 2-AG levels in the vHC exhibited a positive correlation with the intensity of fear generalization. Moreover, susceptibility to trauma-induced behavioral alterations is associated with marked neuronal hypoactivity in the circuitry involved in the regulation of fear responses, alongside altered expression profiles of genes encoding proteins involved in the modulation of neuroplasticity and eCB signaling. Specifically, these alterations were observed in genes governing eCB metabolism enzymes (*Daglb*, *Faah*, *Abhd12*), receptors (*Grm5*), and neuronal cell surface proteins (*Nlgn3*). Employing unsupervised and semi-supervised machine learning-based statistical approaches, we found that hippocampal gene expression patterns have marked predictive power regarding susceptibility to trauma-induced behavioral alterations.

Our findings underscore the inadequacy of assessing the entire population alone, as it may mask trauma-induced variations in eCB levels in the fear circuitry. Specifically, we have shown that basal levels of 2-AG and AEA in the PrL, IL, vHC, and BLA were unaffected in the short or long-term following a traumatic experience. Though we observed that traumatization resulted in significant long-term contextual fear responses, fear generalization did not differ from the control group, and previous shock exposure did not affect 2-AG or AEA levels in the aforementioned brain regions. It is crucial to avoid misinterpreting these results as indicative of no impact of fear conditioning on eCB signaling. However, this perspective overlooks the considerable variability observed in fear generalization among shocked subjects, a feature previously shown in trauma-exposed individuals to differentiate people with or without subsequent development of PTSD (Kaczkurkin et al., 2017). Our results reflect this phenomenon indicating the heterogeneity of individual responses to the effects of trauma experience and emphasizing the importance of utilizing a nuanced model capable of distinguishing between resilient and susceptible individuals, an approach gaining more recognition in the field (Colucci et al., 2020; Pascual Cuadrado et al., 2022).

Employing our model of trauma exposure, we were able to generate the necessary variability in trauma-induced generalized fear responses to be able to identify susceptible and resilient individuals. As the levels of generalized fear showed high variability, these subpopulations markedly differ in this behavioral response. In contrast, neither acute fear responses during traumatic stress exposure, nor long-term contextual fear conditioning responses showed differences between the identified subpopulations. As the ratio of susceptible subjects in our sample was determined to reflect the proportion of trauma survivors subsequently diagnosed with PTSD (i.e. susceptible individuals) in the human population (Breslau et al., 1998, 1991), we developed a model in which PTSD susceptibility and all its behavioral and neuronal correlates can be studied with high translational validity.

Surprisingly, we detected no discernible differences in AEA content associated with susceptibility in our model. This finding contrasts with studies reporting lower circulating AEA levels in human PTSD patients compared to both healthy individuals and trauma-exposed healthy controls(Neumeister et al., 2013). To reconcile this discrepancy, it is important to consider that circulating endocannabinoid (eCB) levels may not necessarily reflect brain eCB content following release from neurons. Notably, eCBs undergo rapid reuptake processes (Fowler, 2013). Furthermore, it is pertinent to recognize that alterations in AEA levels are not uniformly associated with PTSD, as evidenced by reports of unaltered AEA levels in PTSD patients (Hill et al., 2013). Thus, while circulating AEA levels may offer insights in certain contexts, their correlation with PTSD pathology and susceptibility remains an area necessitating further exploration and elucidation.

While our findings seemingly contradict studies in laboratory animals of which the majority reported footshock-induced elevated AEA levels, these works studied the effects immediately after footshock exposure (Morena et al., 2014), following a reminder to the traumatic context (Olango et al., 2012) or after extinction training (Gunduz-Cinar et al., 2013). Furthermore, in the human clinical studies AEA levels were assessed in subjects with the complete PTSD symptom scale while we specifically assessed generalized fear responses as they were the base of our separation of susceptible subpopulations. Interestingly, while changes in AEA content could not be detected concerning susceptibility, there was significant overexpression of *Faah*, the gene of fatty acid amid hydrolase (FAAH), the main degrading enzyme of AEA in the PrL of susceptible subjects in comparison to resilient individuals, suggesting that elevated FAAH activity is linked with a tendency to show trauma-induced fear generalization. This finding confirms that FAAH is involved in the manifestation of PTSD-related symptoms as carriers of C385A single nucleotide polymorphism in the *FAAH* gene resulting in less stable FAAH protein with decreased AEA degrading potency (Sipe et al., 2002) showed less intense symptoms (Crombie et al., 2022; Mayo et al., 2020; Spagnolo et al., 2016). Taken together, while our findings regarding no differences between resilient and susceptible subpopulations in brain AEA content are in contrast with literature findings, this is possibly explained by the source and the time point of sampling. Nevertheless, changes in the expression of the *Faah* gene suggest that AEA signaling might be involved in the basis of susceptibility.

In our investigation of 2-AG content and its association with susceptibility, we have demonstrated that the levels of this eCB were elevated in the PrL of resilient subjects compared to non-shocked or susceptible individuals. This is particularly interesting in light of recent work showing that depletion of 2-AG increases fear generalization and neural correlates related to fear generalization within the PrL (Rosas-Vidal et al., 2024). As such, the elevated 2-AG signaling we found within the PrL of resilient rats, may enhance neural specificity to fear cues and restrict the development of generalization to promote resiliency. In the vHC, we observed a significant experimental group effect on 2-AG content and a strong, albeit statistically non-significant, trend of lower levels in resilient individuals. Furthermore, we discovered a robust positive correlation between vHC 2-AG content and the strength of generalized fear, suggesting that 2-AG signaling in the vHC is indeed associated with susceptibility to develop trauma-induced generalized fear responses.. This relationship is in opposition to the PrL, reinforcing the importance of examining regional specificity in the manner through which eCB signaling may influence fear processing. This finding is in line with human clinical studies have reported correlations between PTSD symptom severity and circulating eCB content (Bluett et al., 2014; Hill et al., 2013). A possible explanation for the elevated vHC 2-AG levels in susceptible individuals could be the elevated hippocampal expression of *Daglb*, the gene encoding diacylglycerol lipase beta (DAGLβ), one of the synthesizing enzymes of 2-AG. Interestingly, the other 2-AG synthesizing enzyme. diacylglycerol lipase alpha (DAGLα), is also directly linked to stress resilience. Conditional deletion of the *Dagla* in vHC has been shown to increase resilience (Kondev et al., 2023). Our finding might be intriguing as DAGLβ has long been proposed not to be involved in the retrograde 2-AG signaling route suppressing synaptic transmission (Gao et al., 2010; Tanimura et al., 2010; Yoshino et al., 2011). However, there is evidence, at least in specific cases, that it indeed contributes to depolarization-induced suppression of excitation mediated by 2-AG (Jain et al., 2013). Moreover, *Daglb* expression is reactive to stress, further validating our findings (Marco et al., 2014).

Intriguingly, in both PrL and vHC, 2-AG levels exhibited a trauma-related pattern. Non-shocked controls and susceptible individuals showed similar 2-AG content while resilient subjects displayed either elevated or decreased levels in a brain site-dependent manner when compared to these groups. While such a pattern might seem difficult to interpret in terms of susceptibility, as 2-AG levels do not necessarily align with the levels of generalized fear across groups, similar phenomena have been observed regarding susceptibility to stress. For instance, both susceptible and non-stressed animals show similar expression levels of sphingosine-1-phosphate receptor 3 in the mPFC following chronic social defeat stress, while resilient animals show elevated levels (Corbett et al., 2019). There are several possible explanations for such a seemingly discrepant pattern. Firstly, 2-AG levels might be differentially involved in the regulation of behavioral responses under physiological and stressed conditions, resulting in similar behavioral phenotypes with different eCB backgrounds. Secondly, susceptible individuals might exhibit a deficit in the neuronal regulation of emotional response to a traumatic experience, which cannot suppress further generalized fear response. Lastly, it is important to note that the direction of the difference found in susceptible compared to resilient subjects is dependent on the brain region, suggesting that the compensatory eCB signaling necessary for healthy behavior observed in resilient subjects has a more general deficit in susceptible individuals throughout the fear circuitry.

Our unsupervised and semi-supervised machine learning-based statistical analyses of gene expression data in resilient and susceptible subjects have underscored the substantial role of the HC in susceptibility. Both approaches confirmed that gene expression patterns in this region possess superior predictive power compared to other assessed areas. Notably, the HC expression of *Npas4*, the gene of transcription factor neuronal PAS domain protein 4 (Npas4). a modulator of neuronal plasticity and marker of neuronal activity, had a predictive power four times greater than any other assessed gene. This gene exhibited a marked downregulation in the susceptible compared to resilient subpopulations across all assessed areas, particularly in the HC. Such downregulation of *Npas4* suggests HC hypoactivity, closely linked to susceptibility. This finding aligns with human clinical data showing that lower HC activation can predict PTSD symptom severity in patients (van Rooij et al., 2018). Furthermore. *Npas4* expression in the HC was shown to be crucial in context coding in a rodent model of PTSD (Weng et al., 2018), suggesting that downregulation of the gene might lead to a deficit in context coding and contribute to the overgeneralization of fearful responses in contexts not associated with the previous traumatic experience. Indeed. *Npas4* downregulation in several brain regions in the fear circuitry was shown to be associated with stress-related psychopathologies (Selçuk et al., 2023). Decreased expression of *Arc*, another immediate early gene that similarly to *Npas4* showed downregulation in our model, was shown to specifically increase susceptibility to trauma-induced adverse behavioral changes (Kozlovsky et al., 2008). Most importantly, Npas4 was shown to directly interact with eCB signaling in the HC, promoting eCB-mediated depolarization-induced suppression of inhibition in the regulation of neuronal plasticity (Hartzell et al., 2018). Taken together, one might hypothesize that a substantial deficit in Npas4-mediated mechanisms linked to changes in eCB signaling fundamentally alters neuronal plasticity, leading to abnormal processing of traumatic experiences, impaired contextual coding, and ultimately, generalized fear responses.

## 5. Conclusions

Our findings reveal that subpopulations differing in their susceptibility to developing PTSD-like symptoms show anatomically specific differences in eCB content. Specifically, we have highlighted that eCB changes in the vHC are crucial for susceptibility, as 2-AG levels in this region showed significant correlations with the strength of generalized fear and are associated with neuronal hypoactivity and decreased expression of crucial mediators of neuronal plasticity. Patterns of gene expression related to neuroplasticity and eCB signaling in the HC had substantial predictive power for PTSD susceptibility. Our results describe a novel neuropharmacological marker that outlines susceptibility to developing PTSD following trauma exposure and may contribute to the identification of clinically relevant subpopulations.

## Author contributions

Conceptualization, M.A., L.Sz., E.M., M.T., and M.N.H.; Methodology, L.Sz., M.A.; Investigation, L.Sz., M.A., Gy.Y.B., B.T., L.B., and Z.B.; Data Curation, L.Sz.; Formal Analysis, L.Sz., V.Z.K.; Writing – Original Draft Preparation, M.A., L.Sz.; Writing – Review & Editing Preparation, L.Sz., Gy.Y.B., Z.K.V., B.T., L.B., Z.B., M.N.H., M.T., E.M, and M.A.; Visualization Preparation, L.Sz., M.A., and V.Z.K.; Supervision, M.A. and E.M.; Project Administration, M.A.; Funding Acquisition, M.A., E.M., M.T., and M.N.H.

## Acknowledgments

We thank the Behavioral Studies Unit core facility of the Institute of Experimental Medicine Hungarian Research Network and the technical assistance of D. Ömböliné and B. Molnar.

## Supplemental information legends

**Supplemental Figure 1. Endocannabinoid levels show no trauma-induced changes in the fear circuitry when assessed in the whole sample regardless of susceptibility. A**) Experimental design schematic showing our model of trauma exposure. Following contextual fear conditioning (Day 0), contextual fear memory expression was assessed 28 days later in the traumatic context (*Context A*. Day 28). The next day, fear generalization was studied in a novel, safe context (*Context B*. Day 29). Subjects were immediately sacrificed after *Context B* testing, and brain samples from prelimbic cortex (PrL), infralimbic cortex (IL), ventral hippocampus (vHC) and basolateral amygdala (BLA) were collected for HPLC/MS/MS analysis of 2-arachidonoylglycerol (2-AG) and N-arachidonoylethanolamin (AEA) levels. Non-shocked controls underwent the same procedure without the delivery of footshocks. **B**) Contextual fear conditioning induced freezing behavior during the conditioning session. (**C**) which remained high during Context A contextual reminder. **D**) The shocked group showed no difference in fear generalization compared to non-shocked control groups. **E-H**) No differences were detected in eCB levels at the assessed areas between shocked and non-shocked groups. ***: p < 0.001; Data are presented as mean ± SEM. One-way ANOVA.

## Abbreviations

2-AG: 2-arachidonoylglycerol
AEA: N-arachidonoylethanolamine, anandamide
AMY: amygdala
BLA: basolateral amygdala
CB1R: cannabinoid receptor type-1
DAGLα: diacylglycerol lipase alpha
DAGLβ: diacylglycerol lipase beta
eCB: endocannabinoid
HC: hippocampus
IL: infralimbic cortex
mPFC: medial prefrontal cortex
Npas4: neuronal PAS domain protein 4
PrL: prelimbic cortex
PTSD: posttraumatic stress disorder
vHC: ventral hippocampus

