## supplementary material for "Endocannabinoid and neuroplasticity-related changes as susceptibility factors in a rat model of posttraumatic stress disorder"

**Supplementary Table 1. Statistical data from the analysis of *Experiment 2* with comparisons of shocked vs. non-shocked group regardless of susceptibility.** 2-AG: 2-arachidonoylglycerol; AEA: N-arachidonoylethanolamine; PrL: prelimbic cortex; IL: infralimbic cortex; vHC: ventral hippocampus; BLA: basolateral amygdala; *: significant group effect. Mann-Whitney U test.

| | **test / area** | **U** | **p-value** |
|---|---|---|---|
| freezing duration | **conditioning** | **0** | **<0.0001*** |
| | ***Context A*** | **0** | **<0.0001*** |
| | *Context B* | 18 | 0.12 |
| 2-AG | PrL | *23.5* | 0.26 |
| | IL | *28* | 0.46 |
| | vHC | *37* | 0.97 |
| | BLA | *34* | 0.78 |
| AEA | PrL | *22.5* | 0.23 |
| | IL | *36* | 0.91 |
| | vHC | *27.5* | 0.42 |
| | BLA | *19.5* | 0.14 |

**Supplementary Table 2. Genes assessed in *Experiment 3*.** 2-AG: 2-arachidonoylglycerol; AEA: N-arachidonoylethanolamine.

| function | gene | protein |
|---|---|---|
| **receptors** | *Cnr1* | Cannabinoid receptor type 1 |
| | *Cnr2* | Cannabinoid receptor type 2 |
| | *Dag1* | Dystroglycan 1 |
| | *Gpr3* | G protein-coupled receptor 3 |
| | *Gpr6* | G protein-coupled receptor 6 |
| | *Gpr12* | G protein-coupled receptor 12 |
| | *Gpr55* | G protein-coupled receptor 55 |
| | *Grm5* | Glutamate metabotropic receptor 5 |
| | *PPara* | Peroxisome proliferator activated receptor alpha |
| | *Slc17a8* | Solute carrier family 17 member 8 |
| | *Trpv1* | Transient receptor potential cation channel. subfamily V. member 1 |
| **signaling** | *Adam17* | ADAM metallopeptidase domain 17 |
| | *Cacna1a* | Calcium voltage-gated channel subunit alpha1 A |
| | *Cacna1b* | Calcium voltage-gated channel subunit alpha1 B |
| | *Cnrip1* | Cannabinoid receptor interacting protein 1 |
| | *Gnai1* | G protein subunit alpha i2 |
| | *Gnai2* | G protein subunit alpha i2 |
| | *Mmp9* | *Matrix metallopeptidase 9* |
| | *Nlgn3* | Neuroligin 3 |
| | *Nrxn1* | Neurexin 1 |
| **AEA metabolism** | *Alox5* | Arachidonate 5-lipoxygenase |
| | *Alox15* | Arachidonate 15-lipoxygenase |
| | *Faah* | Fatty acid amide hydrolase |
| | *Napepld* | N-acyl phosphatidylethanolamine phospholipase D |
| | *Nat1* | N-acetyltransferase 1 |
| | *Plcb1* | Phospholipase C beta 1 |
| | *Ptgs2* | Prostaglandin-endoperoxide synthase 2 |
| | *Ptpn22* | Protein tyrosine phosphatase. non-receptor type 22 |
| **2-AG metabolism** | *Abhd2* | Abhydrolase domain containing 2. acylglycerol lipase |
| | *Abhd6* | Abhydrolase domain containing 6. acylglycerol lipase |
| | *Abhd12* | Abhydrolase domain containing 12. lysophospholipase |
| | *Dagla* | Diacylglycerol lipase. alpha |
| | *Daglb* | Diacylglycerol lipase. beta |
| | *Homer1* | Homer scaffold protein 1 |
| | *Homer2* | Homer scaffold protein 2 |
| | *Mgll* | Monoglyceride lipase |
| | *Plcg1* | Phospholipase C. gamma 1 |
| **transport** | *Fabp5* | Fatty acid binding protein 5 |
| | *Nucb1* | Nucleoindin 1 |
| **activity/neuroplasticity** | *Arc* | Activity-regulated cytoskeleton-associated protein |
| | *Egr1* | Early growth response 1 |
| | *Fos* | Fos proto-oncogene. AP-1 transcription factor subunit |
| | *Fosb* | FosB proto-oncogene. AP-1 transcription factor subunit |
| | *Npas4* | Neuronal PAS domain protein 4 |
| | *Nptx2* | Neuronal pentraxin 2 |
| **housekeeping** | *Actb* | Actin. beta |
| | *Gapdh* | Glyceraldehyde-3-phosphate dehydrogenase |
| | *Rn18s* | 18S ribosomal RNA |
