## Supplementary figures and images for "Endocannabinoid and neuroplasticity-related changes as susceptibility factors in a rat model of posttraumatic stress disorder"

### supplementary figure

#### Supplemental Figure 1

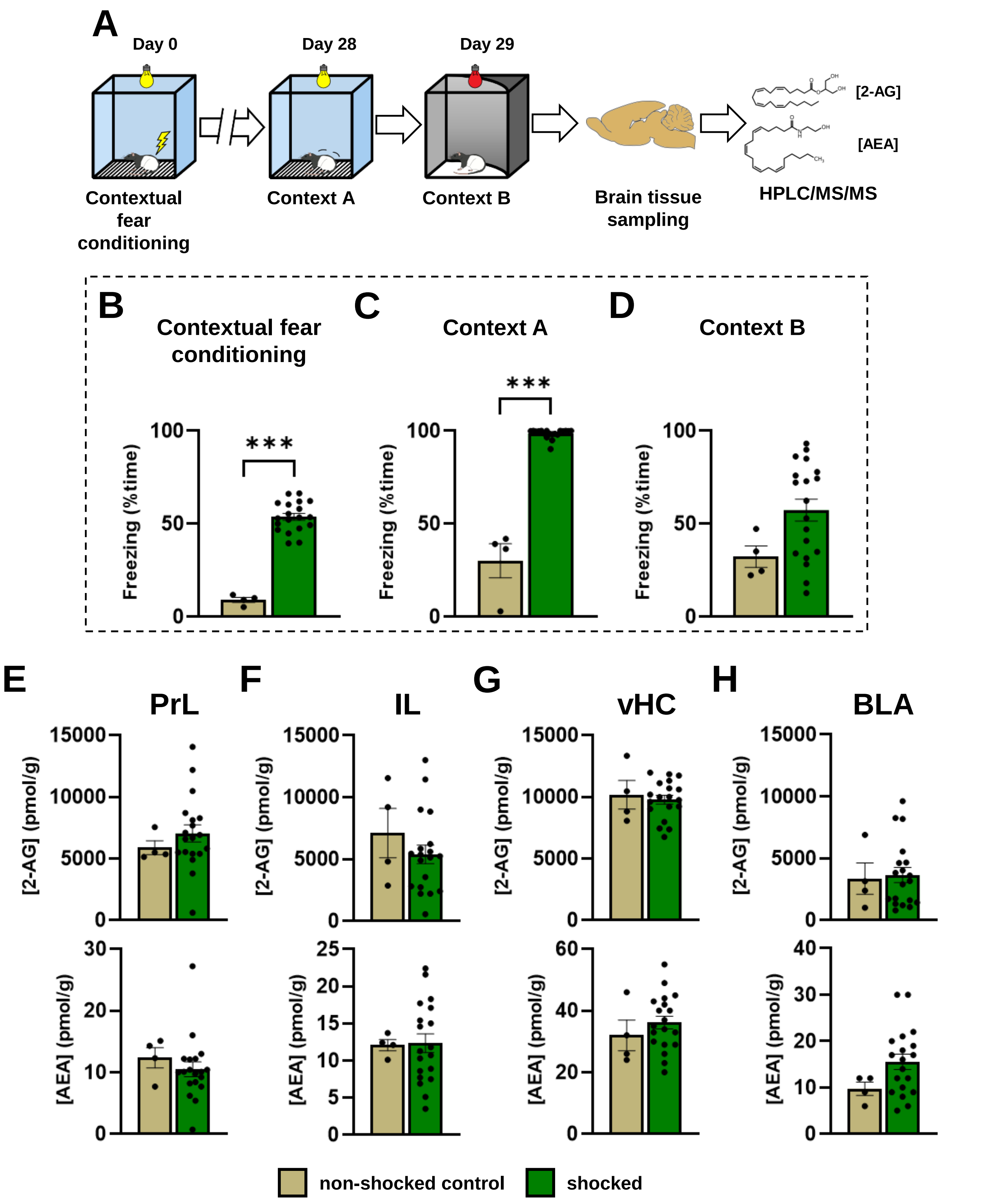
